# Physicochemical Principles Driving Small Molecule Binding to RNA

**DOI:** 10.1101/2024.01.31.578268

**Authors:** Timothy E. H. Allen, James L. McDonagh, Malgorzata Broncel, Carson J. Bryant, Danny Incarnato, Anil Vasudevan, Rabia T. Khan

## Abstract

The possibility of using RNA-targeting small molecules to treat diseases is gaining traction as the next frontier of drug discovery and development. The chemical characteristics of small molecules that bind to RNA are still relatively poorly understood, particularly in comparison to protein-targeting small molecules. To fill this gap, we have generated an unprecedented amount of RNA-small molecule binding data, and used it to derive physicochemical rules of thumb that could be used to define areas of chemical space enriched for RNA binders - the Small molecules Targeting RNA (STaR) rules of thumb. These rules have been applied to publicly available RNA-small molecule datasets and found to be largely generalizable. Furthermore, a number of patented RNA-targeting compounds and FDA-approved compounds also pass these rules, as well as key RNA binding approved drug case studies including Risdiplam. We anticipate this work will significantly accelerate the exploration of the RNA-targeted chemical space, towards unlocking RNA’s potential as a small molecule drug target.

**Graphical Abstract:** 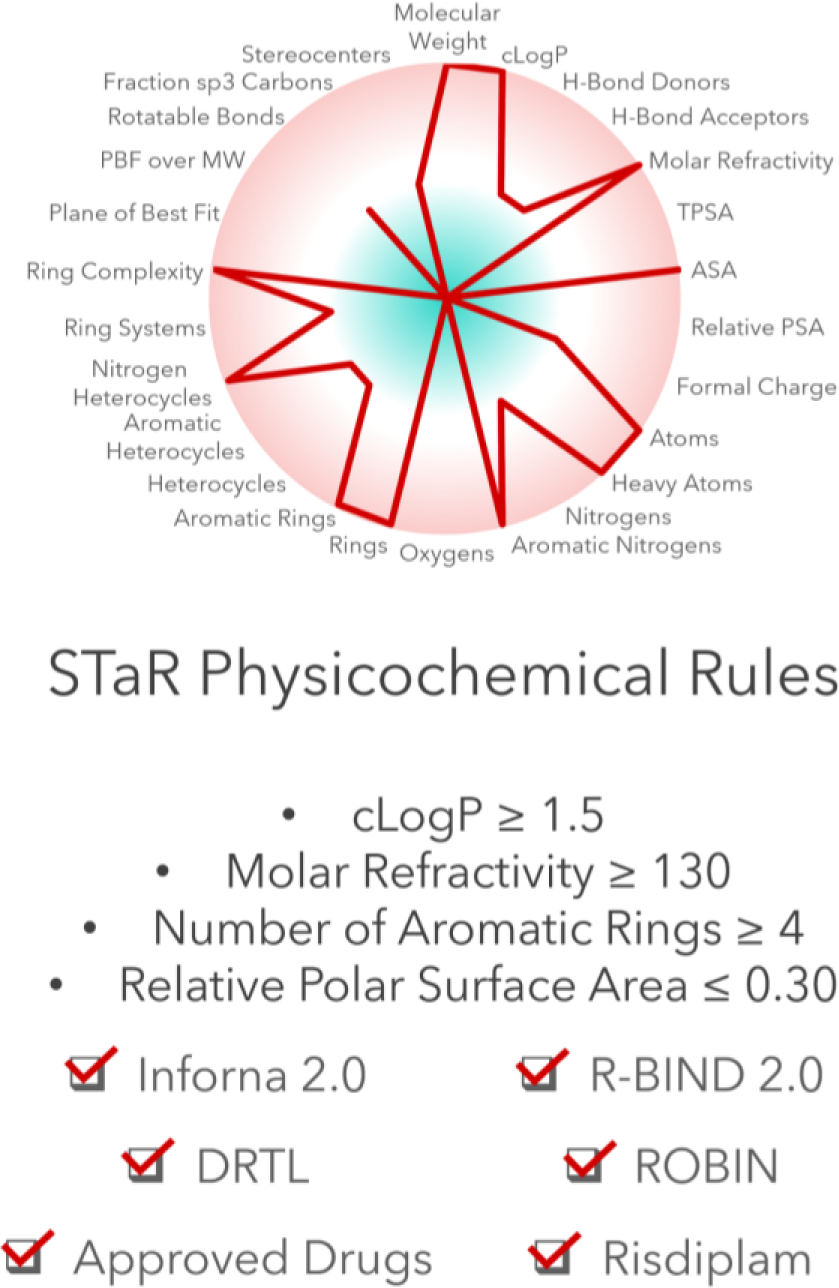

## Introduction

To date, although the industry has referred to the druggable genome, classical drug discovery has focused on the proteome. While approximately 75% of the human genome is transcribed into RNA, ∼1% of the human genome encodes for proteins, and estimates further indicate that less than 15% of the proteome is druggable.^1^ This opens a window of opportunity to target RNA to address unmet medical needs by either modulating previously undruggable proteins by targeting their encoding mRNAs, or non-coding RNA involved in disease onset and progression.

RNA is a compelling target to modulate biology for therapeutic intervention,^2–4^ and targeting RNA with small molecules has gained attention following the approval of Risdiplam in 2020.^5^ RNA is crucial to numerous cellular processes, such as transcription, translation, splicing, and trafficking.^6–8^ Despite the rapid advancements in ASOs, siRNA and novel modalities to target RNA, small molecules represent a compelling therapeutic modality for targeting pockets in RNA structures.^9,10^ Compared to other proposed RNA-targeting modalities like antisense oligonucleotides, small molecules present several advantages including dosing convenience and a well-established development path.^11^ A challenge in developing small molecules that selectively bind to RNA and affect functions such as splicing, is the lack of data that correlates binding to RNA and a functional impact.^12^ Efforts to understand the nature of small molecule-RNA interactions have been led by the Inforna platform,^13,14^ R-BIND^15,16^ and ROBIN,^17^ which is the largest publicly available RNA - small molecule binding data set containing 24,468 unique small molecules.

The published datasets Inforna,^14^ R-BIND,^16^ DRTL^18^ and ROBIN^17^ are publicly available and were analyzed as part of this work. Statistical and modelling comparisons made in R-BIND,^16^ DRTL^18^ and ROBIN^17^ focused on the differences between RNA binders (or RNA functional binders in the cases of R-BIND and DRTL) and FDA-approved drugs, as a surrogate for RNA non-binders. Comparisons to FDA-approved drugs can be a concern as it has been shown that some FDA-approved compounds bind to RNA.^19,20^

### Published Datasets - Inforna

The Inforna^13^ method, pioneered by the Disney group, was originally described in 2014 where it was shown to be capable of identifying active molecules targeting RNA. The method and dataset were later updated in 2016^14^ and contain a total of 244 small molecules, 1331 RNA motifs and 1936 RNA-motif to small molecule interactions. The data is drawn from small molecule microarray experiments performed by the Disney group^13^ and data from a comprehensive scientific literature review identifying RNA binding small molecule data.^14^ The small molecules in the Inforna database were found to be diverse in their chemical structures and to contain privileged RNA-binding chemotypes.^14^ Additionally, the frequency of several small molecule scaffolds was found to be statistically significant in terms of whether a molecule bound to a particular type of RNA motif when compared to all small molecules in the data set.

### Published Datasets - R-BIND and DRTL

R-BIND,^15^ managed by the Hargrove group, is an aggregated dataset of small molecules that bind to RNA and unique to this dataset, these molecules elicit a functional outcome. The Hargrove lab have run multiple analyses, including an extension to the database and chemical property analysis^16^ and the use of R-BIND to create an RNA-focused chemical library, the Duke RNA-Targeted Library (DRTL).^18^ R-BIND collates molecules that show functional effects through RNA interactions and has recently been updated to include 153 unique small molecules, information on 55 RNA targets and around 1,500 data points (Table 2).^16^ In R-BIND, the physicochemical properties of R-BIND small molecules are compared to FDA-approved compounds whose molecular weight was not greater than the highest molecular weight ligand in the R-BIND small molecule dataset, which is used as a proxy for bioactive protein targeting ligands. The authors noted that 18/20 of the properties show statistically significant differences in a Mann-Whitney U test. A summary of these changes is outlined in Table 1.

**Table 1.** A Table outlining statistically significant changes physicochemical properties identified between R-BIND Small Molecules (SMs) and FDA Approved compounds as outlined in Donlic et al. 2022.^16^ Here, statistically significant changes must involve a p-value < 0.05 in a Mann-Whitney U-test

| R-BIND Binders versus FDA Approved | Change for R-BIND binders Versus FDA Approved | P-value |
| --- | --- | --- |
| Molecular Weight | Increasing | <0.001 |
| Number of H-Bond Donors | Increasing | <0.001 |
| Number of H-Bond Acceptors | Increasing | <0.05 |
| LogP | Decreasing | <0.05 |
| Topological Polar Surface Area | Increasing | <0.001 |
| LogD | Decreasing | <0.05 |
| Number of Nitrogens | Increasing | <0.001 |
| Number of Oxygens | Decreasing | <0.001 |
| Number of Rings | Increasing | <0.001 |
| Number of Aromatic Rings | Increasing | <0.001 |
| Number of Heterocycles | Increasing | <0.001 |
| Number of Ring Systems | Increasing | <0.001 |
| Ring Complexity | Increasing | <0.001 |
| Number of Stereocenters | Decreasing | <0.001 |
| Fraction of sp<sup>3</sup> Carbons | Decreasing | <0.001 |
| Van der Waals Surface Area | Increasing | <0.001 |
| Formal Charge | Increasing | <0.001 |
| Accessible Surface Area | Increasing | <0.001 |

**Table 2.**
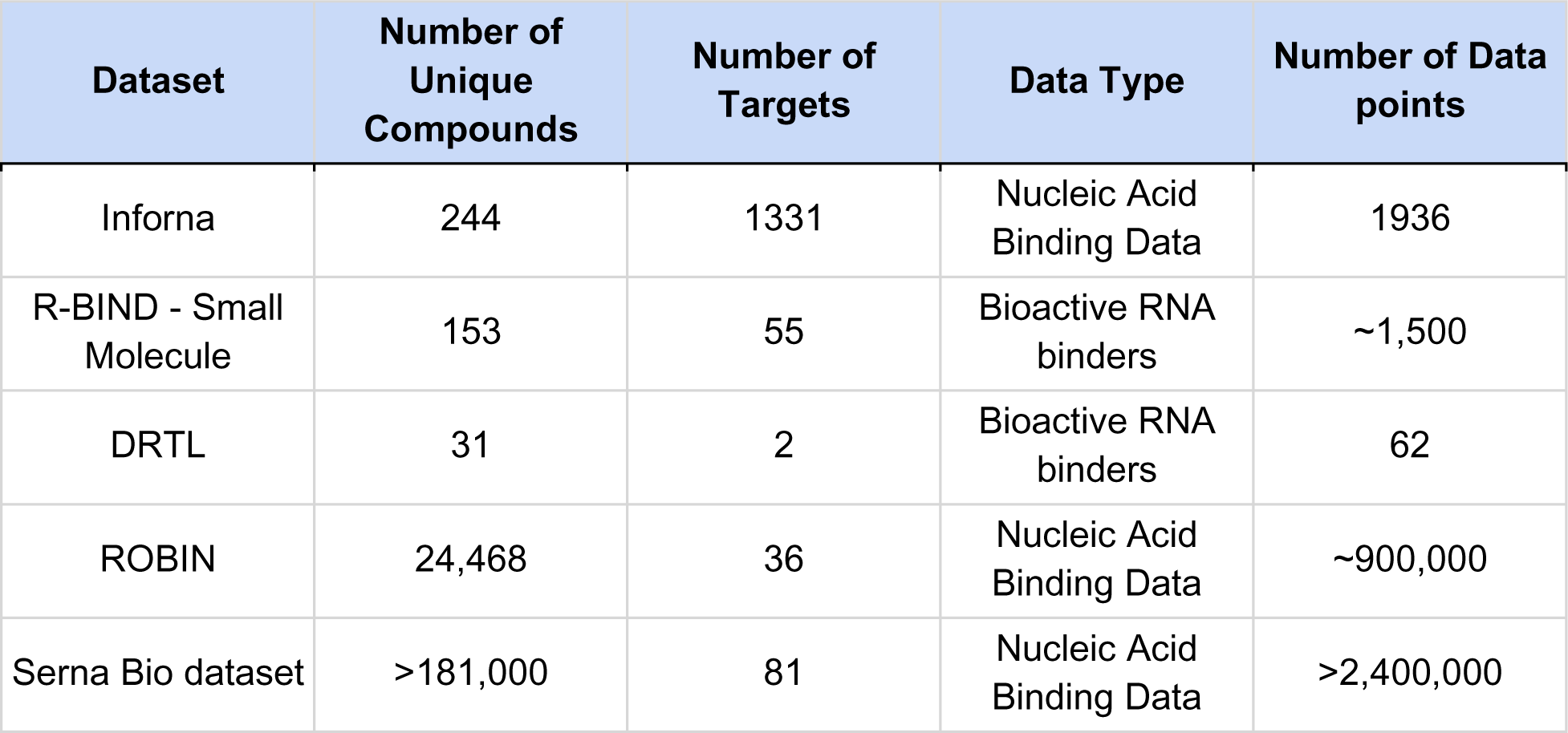
An overview of the datasets examined in this work.

Despite different tendencies in physicochemical properties between RNA bioactive molecules in R-BIND and FDA-approved compounds, Principal Component Analysis has been used to highlight that both sets of molecules display drug-likeness.^16^ The DRTL chemical library constructed from R-BIND using a chemical similarity search was experimentally validated, and data released as part of that study provided 31 Bioactive RNA ligands (Table 2).^18^

### Published Datasets - ROBIN

ROBIN, generated using small molecule microarrays^21^ by the Schneekloth lab,^17^ comprises the largest publicly available dataset of nucleic acid-binding small molecules. This data set is 24,468 unique small molecules screened against 36 nucleic acid targets, including both RNA and DNA biological targets (Table 2), and to our knowledge the largest RNA dataset that includes both binders and non-binders. This data matrix is nearly one million small molecule-nucleic acid interaction data points. Using this data, Kamyar *et al.* developed a classifier capable of differentiating between molecules of the ROBIN RNA binders and the FDA-approved drugs despite a noted overlap in typical medicinal chemistry physicochemical properties such as molecular weight, the number of hydrogen bond donors and acceptors, the number of nitrogens, topological polar surface area and cLogP.

A TMAP reduced dimensionality visualization for chemical space^22^ was able to show that ROBIN RNA binders have diverse chemistry well distributed across chemical space occupied by FDA-approved drugs and protein binders.^17^ The computational work in ROBIN is illustrative of the fact that RNA binders are not easily chemically distinguishable from FDA-approved drugs or protein binders, but that machine learning approaches that can integrate large chemical properties such as the Mordred descriptors,^23^ can help perform this task. The ROBIN dataset is a particularly useful case study for machine learning as it includes data for small molecules that are both binders and non-binders which have been tested against the same set of RNA targets. Supervised machine learning algorithms can make use of this data to train classifiers for the prediction of small molecules as RNA binders or non-binders. The presence of experimental non-binders in ROBIN enables direct comparisons between RNA binders and non-binders that are drawn from the same chemical distribution.

### Published Patented Small Molecules Binding to RNA

As part of this study we have aggregated a list of molecules mined from the patent literature of companies targeting RNA with small molecules, such as Remix Therapeutics, Rgenta Therapeutics and Skyhawk Therapeutics (see the Supplementary Material). While we cannot guarantee these compounds bind to RNA, they are an interesting case study as they have been patented by groups working to develop RNA-targeting therapeutics. This set of data contains 217 unique small molecules.

### In this study

In this work, we aim to extend beyond the work published in R-BIND, DRTL and ROBIN by conducting an unprecedented characterization of RNA binding chemical space using the largest set of experimentally determined RNA small molecule interactions to date. To do this we introduce our proprietary Serna Bio RNA binding dataset (Serna Bio dataset), which comprises more than 2.4 million data points, more than 181,000 unique small molecules, and 81 different RNA targets (Table 2). This dataset includes more than 2,000 approved drug compounds, for which we have analyzed the hit rates to show that approved drugs are more often RNA binders when compared to diverse compound libraries, particularly in the cases of kinase inhibitors, cell cycle and DNA damage repair targeting drugs and epigenetic targeting drugs. Building upon this, we use our dataset to make comparisons between RNA binders and non-binders rather than RNA binders and approved drugs.

A physicochemical analysis has been used to identify key physicochemical properties showing high differentiation between RNA binders and non-binders, specifically cLogP, molar refractivity, the number of aromatic rings and relative polar surface area. These properties have been used to identify physicochemical space enriched for RNA binders, by developing rules of thumb similar to the Rule of Five that has popularly been used to define drug-likeness. The overlap of these rules to druglike chemical space is examined, and suggests that the RNA binder enriched physicochemical space can yield druglike small molecules and that some FDA-approved compounds sit in this space, including the approved drug Risdiplam.^5^ The defined physicochemical space represents an opportunity for exploration in the search for RNA binders and may be useful in developing RNA binders in drug discovery.

## Results and Discussion

### Approved Drugs Bind to RNA

The Serna Bio dataset was collected by performing seven primary screens of commercially available small molecule compound collections, chosen specifically to increase the chemical diversity with each added dataset. These primary biophysical screens were performed using two distinct ASMS technology platforms, the automated ligand identification system (ALIS)^24^ and the self-assembled monolayer desorption ionization-affinity selection-mass spectrometry (SAMDI-ASMS).^25^ These platforms are used to differentiate between RNA binding and non-binding small molecules, and the datasets constructed are similar to ROBIN.^17^ Across these screens, the average hit rate (the percentage of compounds tested at a target found to be binders) for an individual RNA target is 1.2%, which is comparable to the values of 0.2 - 1.2% presented in ROBIN^17^ and 0.01 - 0.9% presented by Rizvi *et al*.^26^

In one primary screen, we found the mean hit rate of FDA-approved drugs to be 1.3% - higher than that for a diverse chemical collection against the same 35 RNA targets (0.8%). Table 3 and Figure 1 show a comparison of the hit rates of different classes of approved drugs against RNA targets to a diverse compound library tested against the same RNA targets.

**Table 3.** a table showing the mean hit rate of various compound collections in the Serna Bio ASMS data - focusing on different classes of approved drug compounds. p-values calculated using Fisher’s Exact test.^27^ p-values found to be statistically significant (α = 0.01) are marked with an asterisk.

| Compound Set | Number of Compounds | Mean Hit Rate (%) | p-value (compared to Diverse Compounds) |
| --- | --- | --- | --- |
| Diverse Compounds | 32451 | 0.8 | n/a |
| FDA Approved Drugs | 1289 | 1.3 | 1.75E-03* |
| Kinase Inhibitors | 98 | 2.7 | 8.58E-03* |
| GPCR-Related | 173 | 1.0 | 1.16E-01 |
| CC-DDR Targeting | 82 | 4.0 | 1.62E-03* |
| Epigenetic Targeting | 33 | 5.0 | 4.85E-04* |
| Antibacterial | 111 | 1.5 | 1.62E-01 |
| Antiviral | 43 | 2.3 | 9.15E-01 |

**Figure 1.**
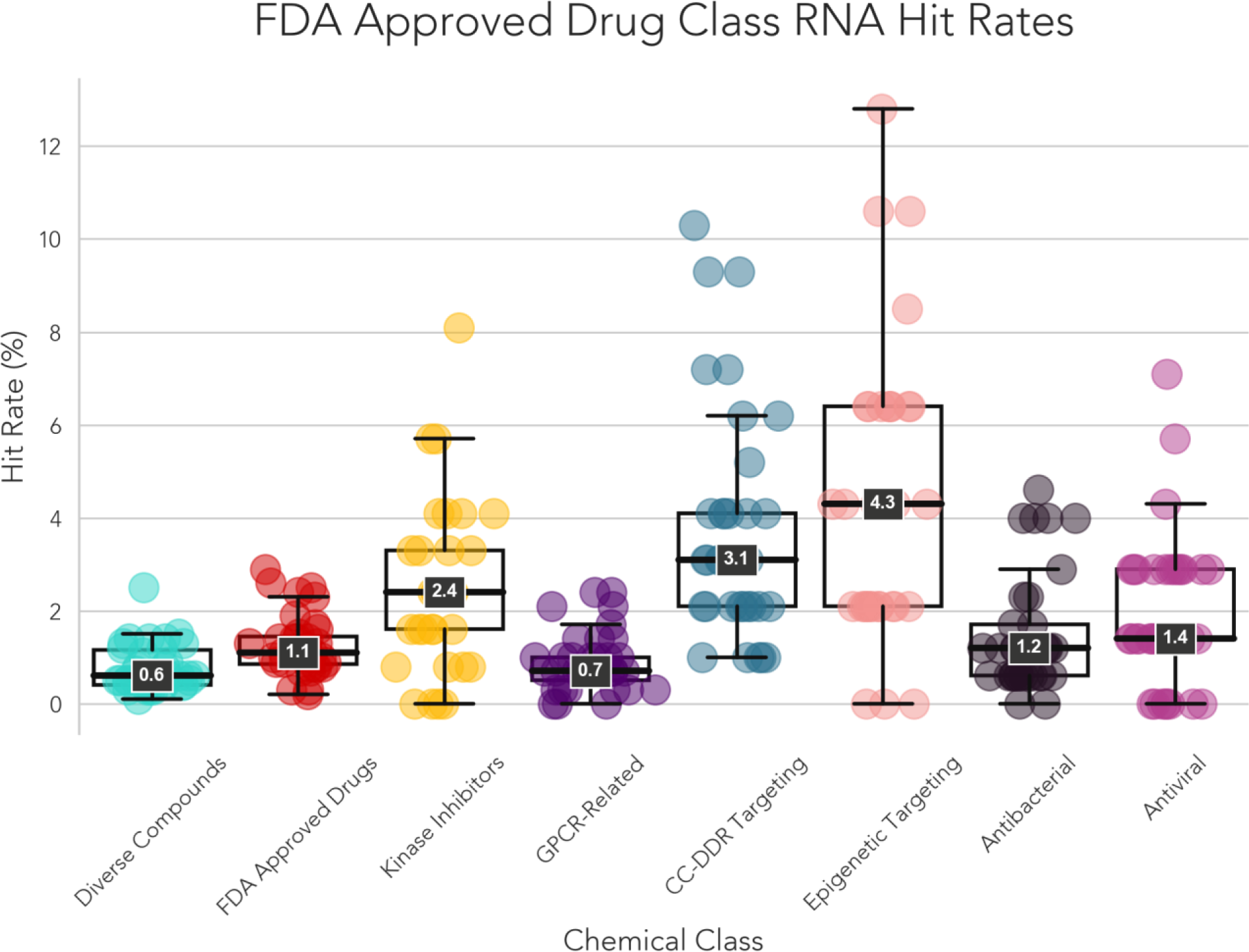
A box and swarm plot showing the hit rate variation between different compound classes each screened against 35 RNA targets. Each point represents the hit rate (the percentage of compounds found to be binders) for that compound class against a specific RNA target. Boxes and whiskers represent the distributions of these hit rates and the median hit rates are shown at the 50th percentile lines.

As published in Fang *et al.*^19^ we also find approved drugs show a statistically significant increase (p < 0.01) in hit rate compared to diverse compounds. The hit rate varies across different classes of approved drugs, with kinase inhibitors, cell cycle and DNA damage repair (CC-DDR) targeting compounds, and epigenetic targeting compounds having the highest hit rates. These findings are aligned with Fang *et al*,^19^ but it should be noted that the concentrations these compounds were tested at are much higher than the physiological concentrations these drugs would typically be dosed at.

As previous work had focused on comparing RNA binders to FDA-approved drugs, we identified the need for a direct comparison of the physicochemical properties of RNA binders to non-binders to provide a greater understanding of the chemical characteristics of RNA binders. It should be noted that RNA binders and non-binders used in this work are classified based on the RNA targets they have been screened against, and non-binders may bind to RNA targets other than those they were tested against.

### Physicochemical Distributions in RNA Datasets

In order to assess if we can define physicochemical properties that differentiate between RNA binders and non-binders, we calculated 26 physicochemical properties (Table 4) for the publicly available RNA-small molecule datasets Inforna 2.0,^13^ R-BIND 2.0,^16^ DRTL^18^ and ROBIN^17^ as well as the Patent data collected for this study and our own in-house Serna Bio dataset, comprising of more than 181,000 unique small molecules tested against at least one of 81 RNA targets across a total of seven primary screens. For the ROBIN and Serna Bio datasets, binders were defined as compounds found to bind to any of the tested RNA targets (details of the experimental protocols are available in the Materials and Methods section), and non-binders were defined as compounds binding to none of the tested RNA targets. For Inforna, R-BIND, DRTL and the Patent data, which contain only binders and no non-binders, a non-binder dataset was constructed using all the non-binders in the Serna Bio and ROBIN datasets and used in the comparison. It should be noted that differences in platform technology, the concentration of tested targets and compounds, and data processing between datasets can lead to differences in hit rates.

**Table 4.**
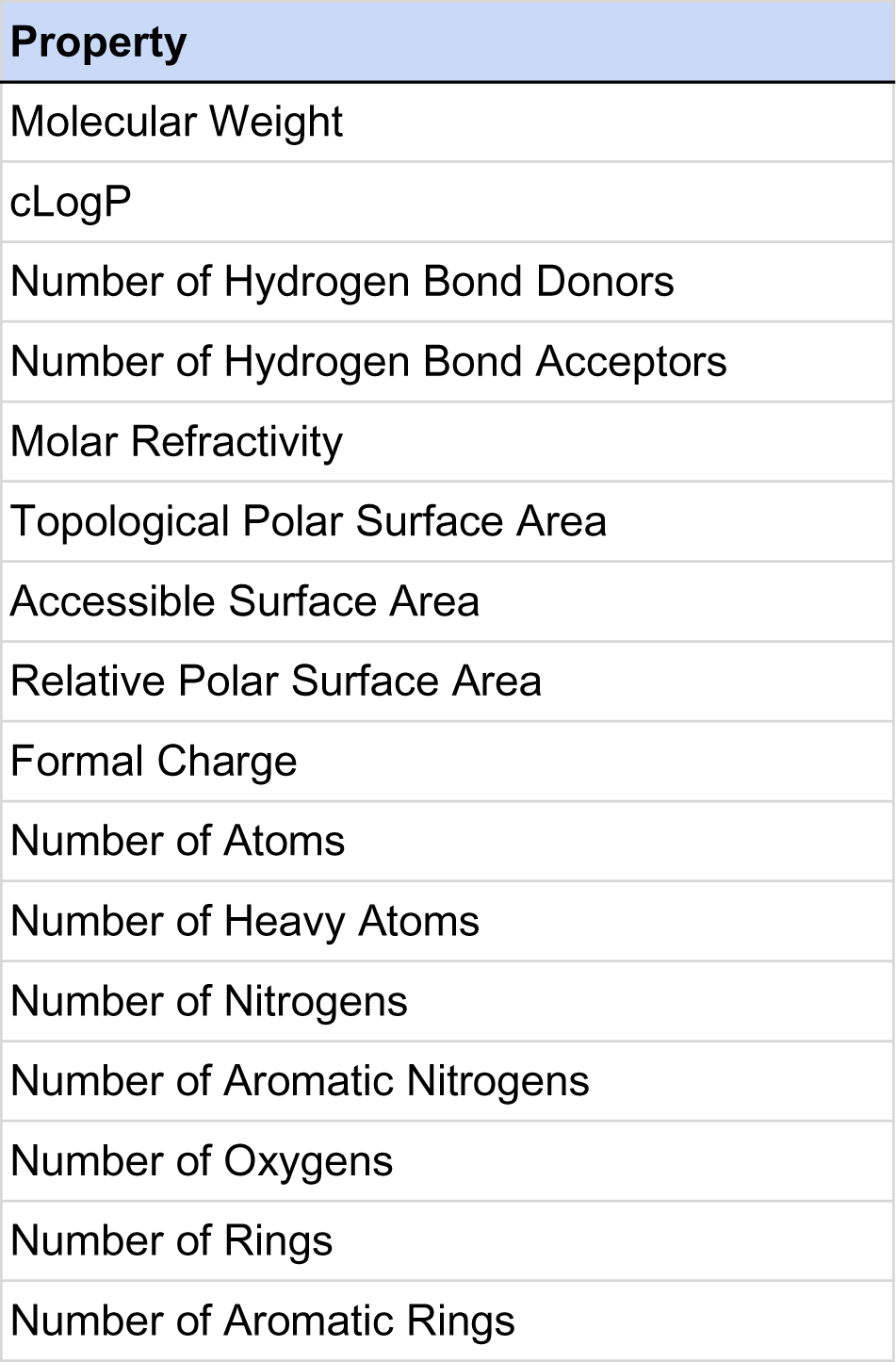
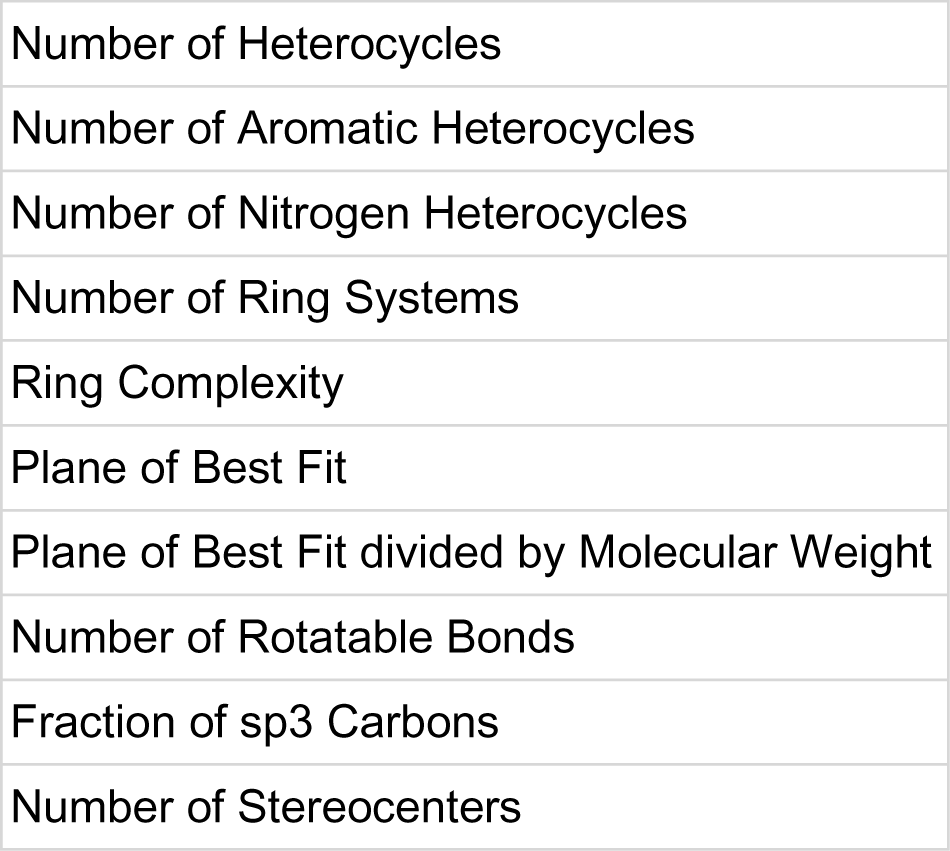
A list of the physicochemical properties calculated in this work.

An overview of the comparison of these physicochemical properties for RNA binders and non-binders is shown in Figure 2 and tabulated in Table 5. Figure 2 shows the physicochemical profiles of RNA binders and non-binders differ quite notably between the different datasets, although there are some commonalities. For example, the relationship between RNA binders and non-binders for the medicinal chemistry properties used in Lipinski’s rules^28^ (the upper right region of the circle) is the same for ROBIN and Serna Bio compounds, illustrated by those areas of the plot having the same shape. Similarity in atomic counts (the lower right region of the circle) is noticed between R-BIND and Serna Bio - increases in the number of atoms, the number of heavy atoms and the number of aromatic nitrogens are common for RNA binders in these datasets. Ring counts (the lower left region of the circle) show similarity between R-BIND, DRTL, Inforna and Serna Bio, with all four datasets identifying RNA binders as having more total rings, more aromatic rings and more nitrogen heterocycles. R-BIND, DRTL and Serna Bio also show RNA binders have greater ring complexity. Finally, R-BIND, ROBIN, Inforna, Serna Bio and the Patent Compounds show commonality in properties measuring compound planarity - the plane of best fit and plane of best fit divided by molecular weight decrease in RNA binders in all these datasets.

**Figure 2.**
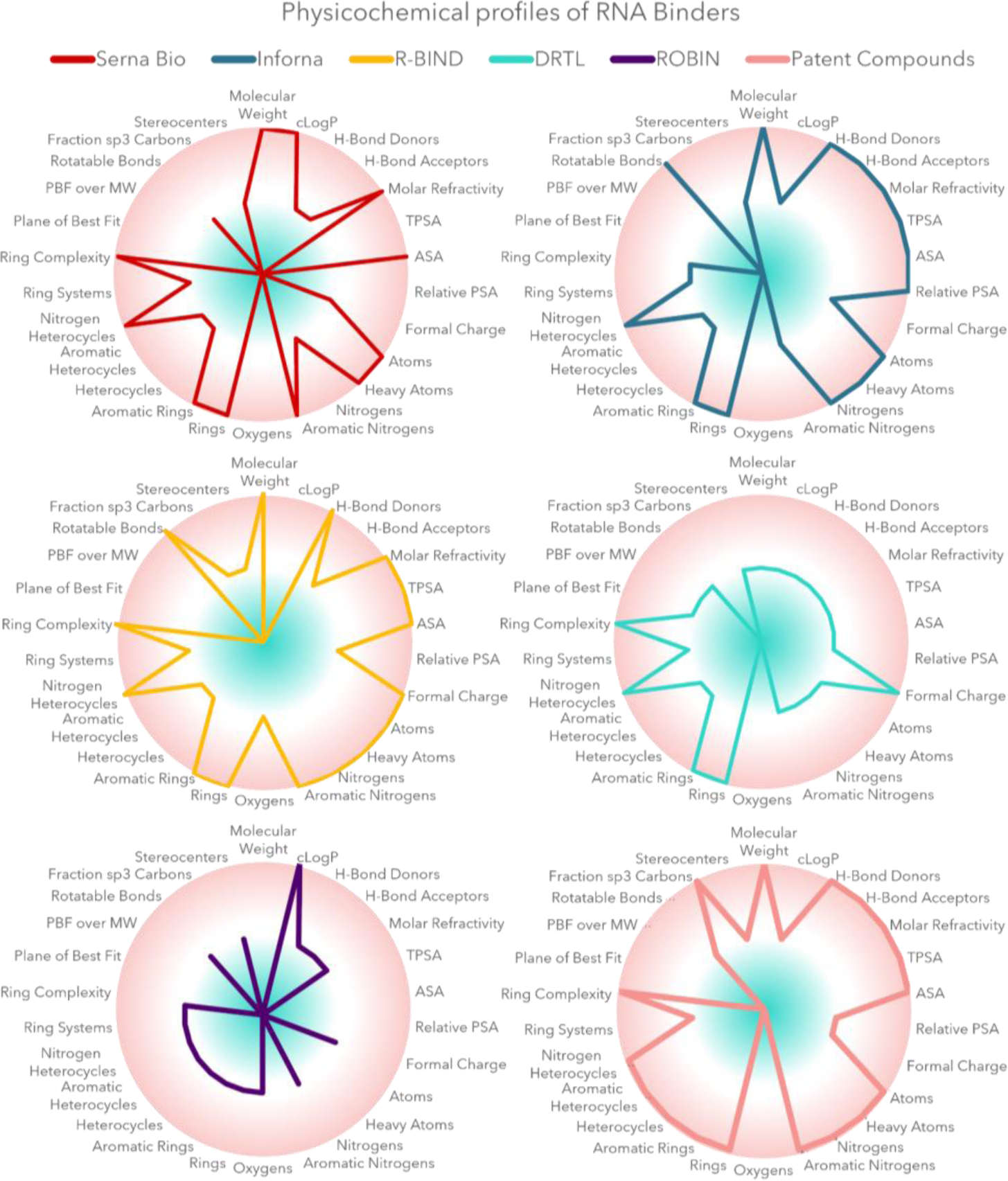
Radar plots showing the change in physicochemical properties between RNA binders and non-binders in the datasets Serna Bio (upper left, red), Inforna (upper right, blue), R-BIND 2.0 (middle left, yellow), DRTL (middle right, turquoise), ROBIN (lower left, purple), and Patent compounds (lower right, pink). The comparison between binders and non-binders in each case is the same as those outlined in Table 5. In these plots, an extension of the line to the edge of the circle indicates a statistically significant increase in that property in RNA binders compared to non-binders, while a contraction to the center indicates a statistically significant decrease in that property in RNA binders compared to non-binders. Here, statistically significant changes must involve a p-value < 0.01 and a change in median value for the physicochemical property in question using Mood’s median test^29^ with Benjamini-Hochberg correction.^30^

**Table 5.**
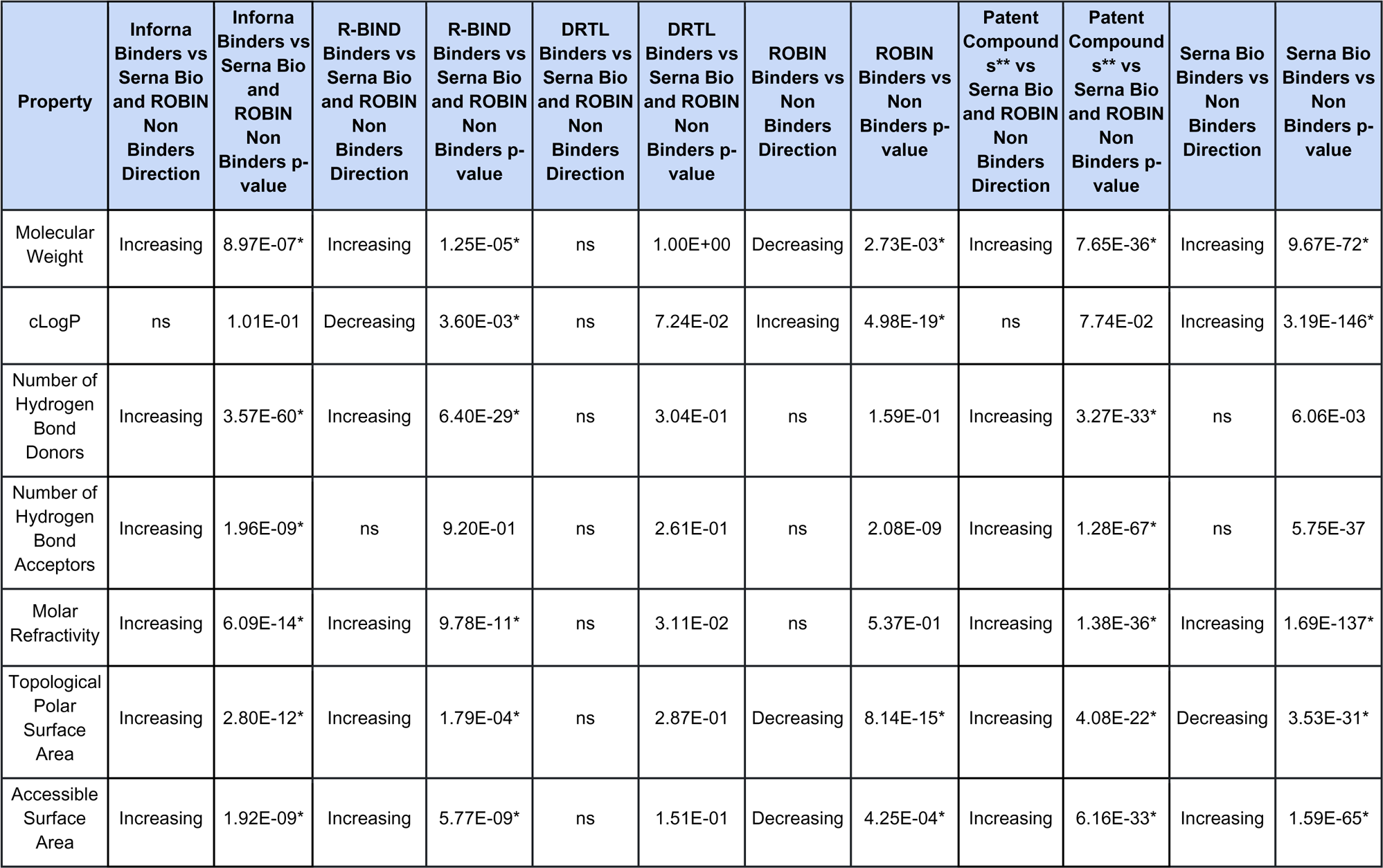
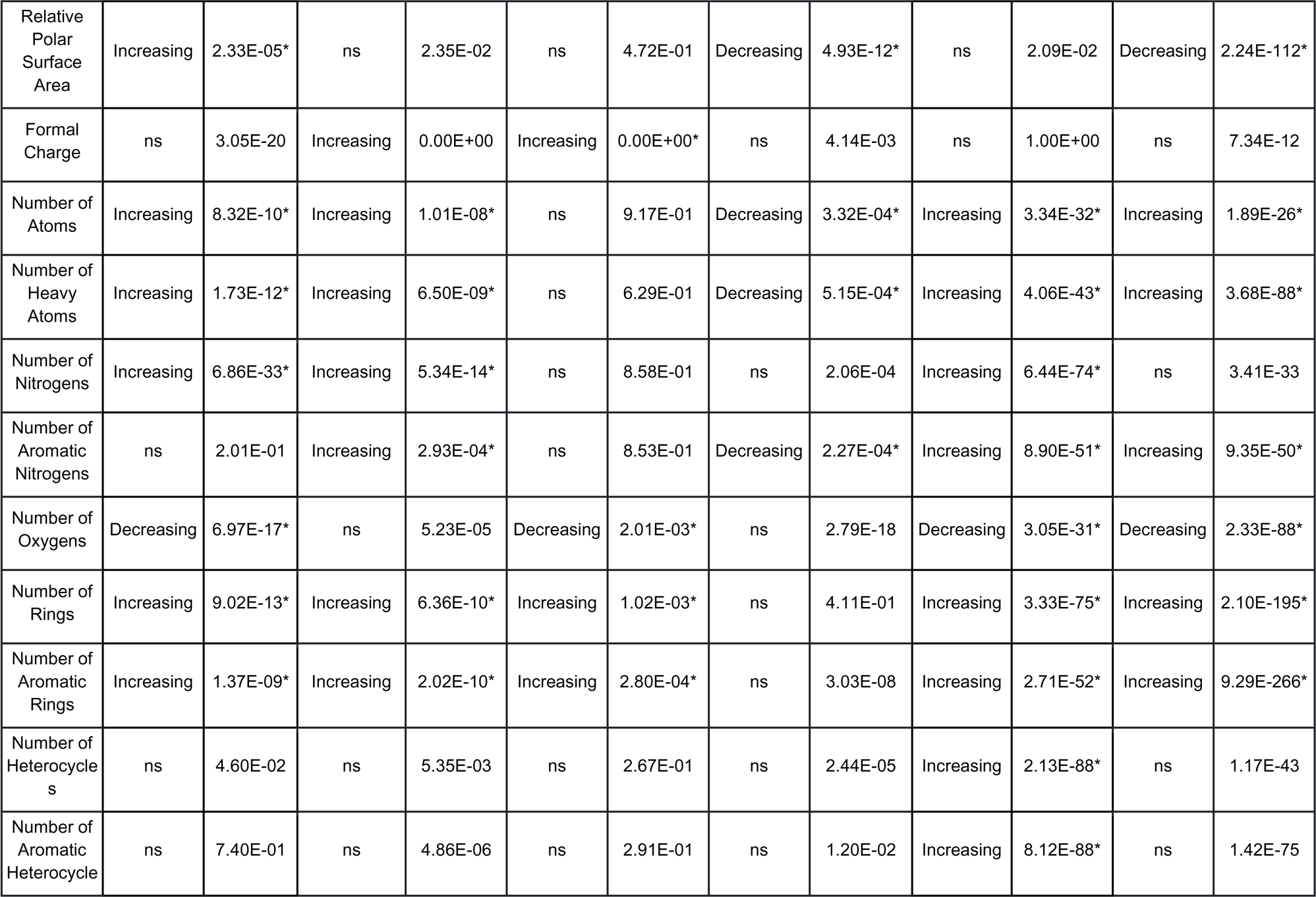
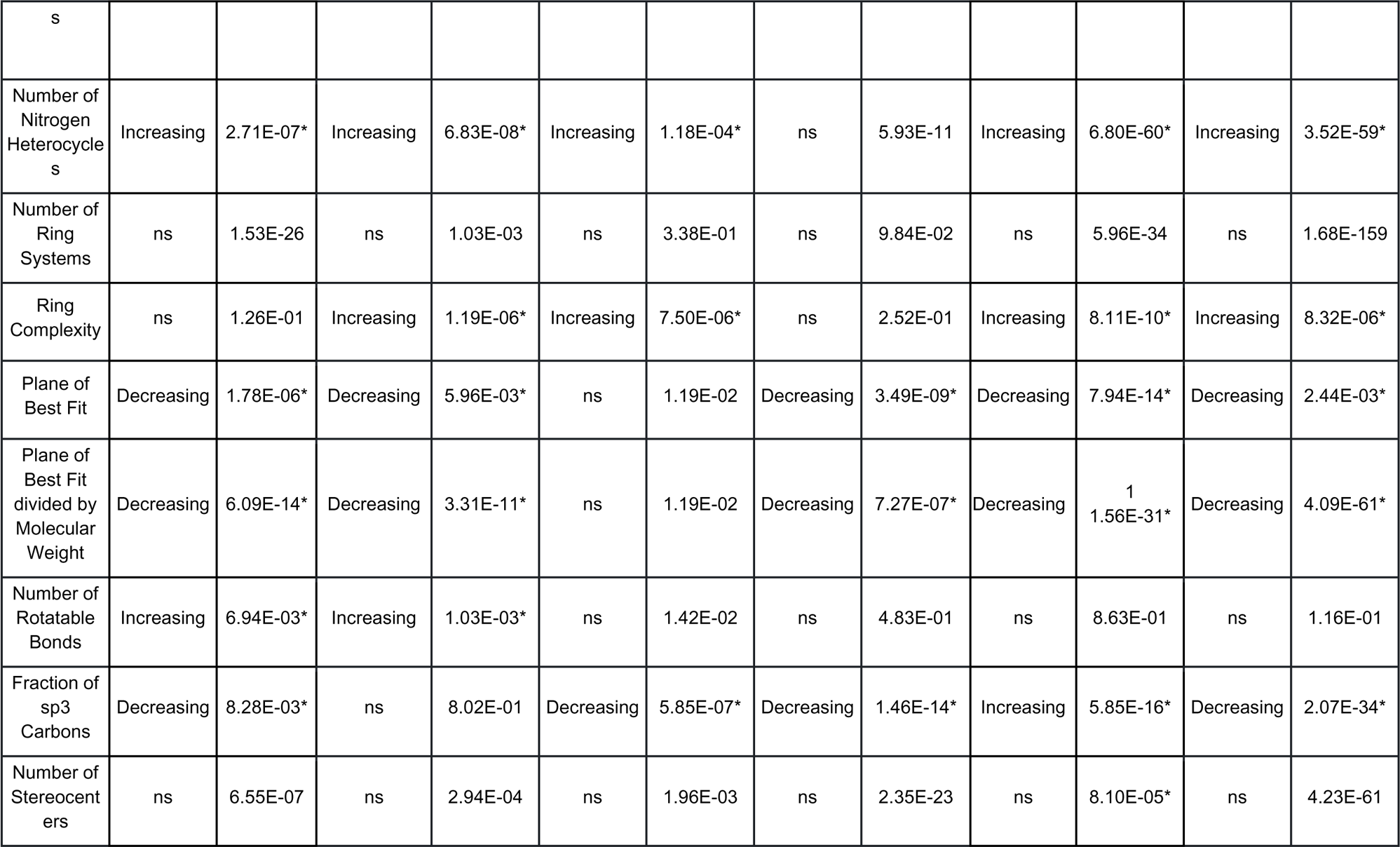
A Table outlining statistically significant changes in physicochemcial properties identified between RNA binders and non-binders for each dataset studied. Here, statistically significant changes must involve a p-value < 0.01 and a change in median value for the physicochemical property in question. Significant changes based on this definition are marked with an asterisk. ns=not significant. Please note **binding data is not confirmed but assumed based upon patent protection.

| Property | Inforna Binders vs Serna Bio and ROBIN Non Binders Direction | Inforna Binders vs Serna Bio and ROBIN Non Binders p-value | R-BIND Binders vs Serna Bio and ROBIN Non Binders Direction | R-BIND Binders vs Serna Bio and ROBIN Non Binders p-value | DRTL Binders vs Serna Bio and ROBIN Non Binders Direction | DRTL Binders vs Serna Bio and ROBIN Non Binders p-value | ROBIN Binders vs Non Binders Direction | ROBIN Binders vs Non Binders p-value | Patent Compound s** vs Serna Bio and ROBIN Non Binders Direction | Patent Compound s** vs Serna Bio and ROBIN Non Binders p-value | Serna Bio Binders vs Non Binders Direction | Serna Bio Binders vs Non Binders p-value |
| --- | --- | --- | --- | --- | --- | --- | --- | --- | --- | --- | --- | --- |
| Molecular Weight | Increasing | 8.97E-07* | Increasing | 1.25E-05* | ns | 1.00E+00 | Decreasing | 2.73E-03* | Increasing | 7.65E-36* | Increasing | 9.67E-72* |
| cLogP | ns | 1.01E-01 | Decreasing | 3.60E-03* | ns | 7.24E-02 | Increasing | 4.98E-19* | ns | 7.74E-02 | Increasing | 3.19E-146* |
| Number of Hydrogen Bond Donors | Increasing | 3.57E-60* | Increasing | 6.40E-29* | ns | 3.04E-01 | ns | 1.59E-01 | Increasing | 3.27E-33* | ns | 6.06E-03 |
| Number of Hydrogen Bond Acceptors | Increasing | 1.96E-09* | ns | 9.20E-01 | ns | 2.61E-01 | ns | 2.08E-09 | Increasing | 1.28E-67* | ns | 5.75E-37 |
| Molar Refractivity | Increasing | 6.09E-14* | Increasing | 9.78E-11* | ns | 3.11E-02 | ns | 5.37E-01 | Increasing | 1.38E-36* | Increasing | 1.69E-137* |
| Topological Polar Surface Area | Increasing | 2.80E-12* | Increasing | 1.79E-04* | ns | 2.87E-01 | Decreasing | 8.14E-15* | Increasing | 4.08E-22* | Decreasing | 3.53E-31* |
| Accessible Surface Area | Increasing | 1.92E-09* | Increasing | 5.77E-09* | ns | 1.51E-01 | Decreasing | 4.25E-04* | Increasing | 6.16E-33* | Increasing | 1.59E-65* |
| Relative Polar Surface Area | Increasing | 2.33E-05* | ns | 2.35E-02 | ns | 4.72E-01 | Decreasing | 4.93E-12* | ns | 2.09E-02 | Decreasing | 2.24E-112* |
| Formal Charge | ns | 3.05E-20 | Increasing | 0.00E+00 | Increasing | 0.00E+00* | ns | 4.14E-03 | ns | 1.00E+00 | ns | 7.34E-12 |
| Number of Atoms | Increasing | 8.32E-10* | Increasing | 1.01E-08* | ns | 9.17E-01 | Decreasing | 3.32E-04* | Increasing | 3.34E-32* | Increasing | 1.89E-26* |
| Number of Heavy Atoms | Increasing | 1.73E-12* | Increasing | 6.50E-09* | ns | 6.29E-01 | Decreasing | 5.15E-04* | Increasing | 4.06E-43* | Increasing | 3.68E-88* |
| Number of Nitrogens | Increasing | 6.86E-33* | Increasing | 5.34E-14* | ns | 8.58E-01 | ns | 2.06E-04 | Increasing | 6.44E-74* | ns | 3.41E-33 |
| Number of Aromatic Nitrogens | ns | 2.01E-01 | Increasing | 2.93E-04* | ns | 8.53E-01 | Decreasing | 2.27E-04* | Increasing | 8.90E-51* | Increasing | 9.35E-50* |
| Number of Oxygens | Decreasing | 6.97E-17* | ns | 5.23E-05 | Decreasing | 2.01E-03* | ns | 2.79E-18 | Decreasing | 3.05E-31* | Decreasing | 2.33E-88* |
| Number of Rings | Increasing | 9.02E-13* | Increasing | 6.36E-10* | Increasing | 1.02E-03* | ns | 4.11E-01 | Increasing | 3.33E-75* | Increasing | 2.10E-195* |
| Number of Aromatic Rings | Increasing | 1.37E-09* | Increasing | 2.02E-10* | Increasing | 2.80E-04* | ns | 3.03E-08 | Increasing | 2.71E-52* | Increasing | 9.29E-266* |
| Number of Heterocycles | ns | 4.60E-02 | ns | 5.35E-03 | ns | 2.67E-01 | ns | 2.44E-05 | Increasing | 2.13E-88* | ns | 1.17E-43 |
| Number of Aromatic Heterocycle | ns | 7.40E-01 | ns | 4.86E-06 | ns | 2.91E-01 | ns | 1.20E-02 | Increasing | 8.12E-88* | ns | 1.42E-75 |
| s |  |  |  |  |  |  |  |  |  |  |  |  |
| Number of Nitrogen Heterocycles | Increasing | 2.71E-07* | Increasing | 6.83E-08* | Increasing | 1.18E-04* | ns | 5.93E-11 | Increasing | 6.80E-60* | Increasing | 3.52E-59* |
| Number of Ring Systems | ns | 1.53E-26 | ns | 1.03E-03 | ns | 3.38E-01 | ns | 9.84E-02 | ns | 5.96E-34 | ns | 1.68E-159 |
| Ring Complexity | ns | 1.26E-01 | Increasing | 1.19E-06* | Increasing | 7.50E-06* | ns | 2.52E-01 | Increasing | 8.11E-10* | Increasing | 8.32E-06* |
| Plane of Best Fit | Decreasing | 1.78E-06* | Decreasing | 5.96E-03* | ns | 1.19E-02 | Decreasing | 3.49E-09* | Decreasing | 7.94E-14* | Decreasing | 2.44E-03* |
| Plane of Best Fit divided by Molecular Weight | Decreasing | 6.09E-14* | Decreasing | 3.31E-11* | ns | 1.19E-02 | Decreasing | 7.27E-07* | Decreasing | <sup>1</sup><br>1.56E-31* | Decreasing | 4.09E-61* |
| Number of Rotatable Bonds | Increasing | 6.94E-03* | Increasing | 1.03E-03* | ns | 1.42E-02 | ns | 4.83E-01 | ns | 8.63E-01 | ns | 1.16E-01 |
| Fraction of sp<sup>3</sup> Carbons | Decreasing | 8.28E-03* | ns | 8.02E-01 | Decreasing | 5.85E-07* | Decreasing | 1.46E-14* | Increasing | 5.85E-16* | Decreasing | 2.07E-34* |
| Number of Stereocenters | ns | 6.55E-07 | ns | 2.94E-04 | ns | 1.96E-03 | ns | 2.35E-23 | ns | 8.10E-05* | ns | 4.23E-61 |

The p-values in Table 5 provide guidance as to which physicochemical properties most effectively differentiate between RNA binders and non-binders for each dataset. Those with the lowest p-values in the Serna Bio binders versus non-binders comparison were considered to construct physicochemical rules to define chemical space that could be investigated for enrichment in RNA binders. In this case that translates to:

- cLogP (being generally greater in RNA binders)
- Molar Refractivity (being generally greater in RNA binders)
- The Number of Aromatic Rings (being generally greater in RNA binders)
- The Relative Polar Surface Area (being generally lower in RNA binders)

All these physicochemical trends are also shared by at least one of the other datasets. The number of rings was not added despite having a low p-value as it was considered redundant to include both the number of rings and the number of aromatic rings, and the p-value is lower in the case of the latter. The choice of using the Serna Bio dataset as the primary source for these property choices was due to the larger number of RNA binders and compounds in total in this dataset.

### Defining RNA Binder-Enriched Physicochemical Space

We set out to identify a Small molecules Targeting RNA (STaR) ruleset that would account for more than 50% of the Serna Bio RNA binders while returning a higher proportion of RNA binders compared to non-binders. This data set was chosen to build the STaR rules as it was the largest, with the other sets being used to evaluate the STaR rule generalizability. The full list of calculated rulesets can be seen in the Supplementary Material. The best-identified ruleset was then converted into the STaR ‘rules of thumb’, which state that, in general, RNA binders have at least two of the following four properties:

○ cLogP ≥ 1.5
○ Molar Refractivity ≥ 130
○ Number of Aromatic Rings ≥ 4
○ Relative Polar Surface Area ≤ 0.30

The STaR rules of thumb were then applied to Serna Bio, R-BIND 2.0, DRTL and ROBIN (Table 6). These results can also be summarized regarding the number of binders and non-binders in each dataset that adhere to the identified STaR rules:

- When identifying compounds in the Serna Bio dataset the STaR rules account for 51% of the RNA binders, 29% of the non-binders and 30% of the total compounds (of all compounds in the dataset)
- When identifying compounds in the ROBIN dataset the STaR rules account for 38% of the RNA binders, 28% of the non-binders and 29% of the total compounds
- When identifying compounds in the R-BIND dataset the STaR rules account for 46% of the RNA binders (there are no non-binders given in the R-BIND dataset)
- When identifying compounds in the DRTL dataset the STaR rules account for 65% of the RNA binders (there are no non-binders given in the DRTL dataset)
- When identifying compounds in the Inforna dataset the STaR rules account for 51% of the RNA binders (there are no non-binders given in the Inforna dataset)
- When identifying compounds in the Patent Data the STaR rules account for 41% of the molecules

**Table 6.** A table showing the results when applying the STaR Rules for RNA binders to Serna Bio, ROBIN, Inforna, Patent Data, R-BIND and DRTL. The Compound Hit Rate and Percentage of non-binding molecules meeting the STaR rules for R-BIND, Inforna, patent data and DRTL is n/a because these datasets are comprised exclusively of RNA binders. Compound hit rate is defined as the ratio of the number of binders correctly identified over the number of molecules identified by the STaR rules. Please note *binding data is not confirmed but assumed based upon patent protection.

| Dataset | Number of Compounds | Number of Binders | Compound Hit Rate | Percentage of binding molecules meeting the STaR rules | Percentage of non-binding molecules meeting the STaR rules | Percentage of all molecules meeting the STaR rules |
| --- | --- | --- | --- | --- | --- | --- |
| Serna Bio | 54868 | 2832 | 0.05 | 51 | 29 | 30 |
| R-BIND | 70 | 70 | n/a | 46 | n/a | 46 |
| DRTL | 20 | 20 | n/a | 65 | n/a | 65 |
| ROBIN | 7038 | 729 | 0.10 | 38 | 28 | 29 |
| Inforna | 109 | 109 | n/a | 51 | n/a | 51 |
| Patent Data | 89 | 89* | n/a | 41 | n/a | 41 |

In the cases of the Serna Bio and ROBIN datasets, the STaR rules identify a higher proportion of RNA binders than non-binders, and the percentage of R-BIND, DRTL and Inforna RNA binders that are identified by the STaR rules are also high. These results indicate a certain level of extensibility of the STaR rules to other RNA binding datasets and support the hypothesis that they are enriching for RNA binder chemical space.

The STaR rules we have outlined above can be visualized in kernel density estimate distribution plots, and these are shown in Figures 3-6. Each plot shows the distribution of RNA binders in the R-BIND, DRTL, ROBIN and Serna Bio datasets, and FDA Approved Compounds,^31^ with the shaded area indicating areas falling outside that specific STaR rule. In terms of the distribution of properties shown in Figures 3-6, it appears as if the majority of compounds pass the cLogP rule, with the other STaR rules then being more selective. As compounds are required to only meet two of the STaR rules to pass, compounds are able to fail on some and still fit inside our RNA binder-enriched area of physicochemical space. A further comparison of Serna Bio and ROBIN binders and non-binders is included in the Supplementary Material (Figures S1-S4).

**Figure 3.**
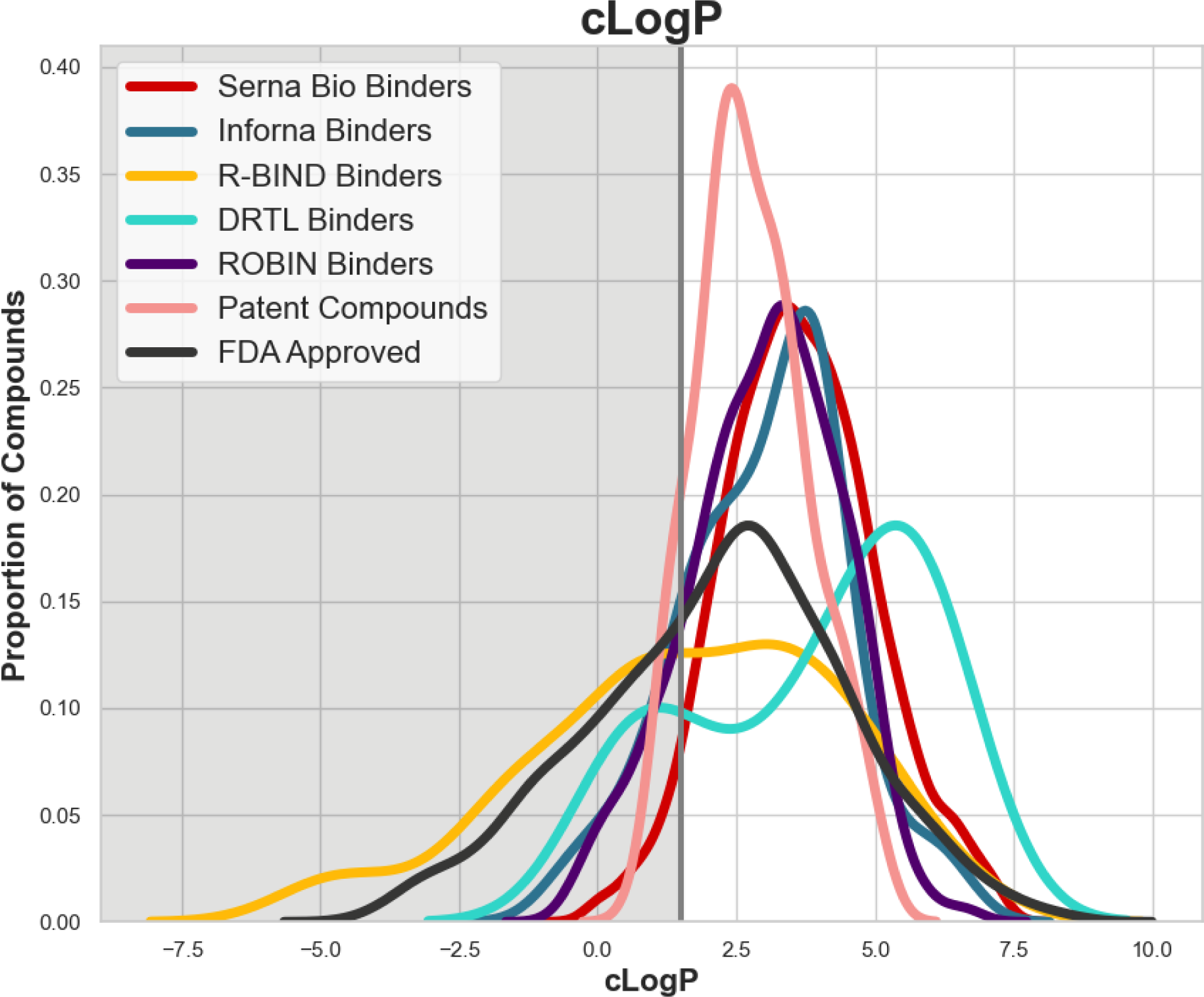
A kernel density estimate distribution plot showing the distribution of cLogP values for Serna Bio binders (red), ROBIN binders (purple), R-BIND binders (yellow), DRTL binders (turquoise), Patent Compounds (pink), Inforna Binders (blue) and FDA Approved Drugs (grey) with a comparison to the rule we have outlined concerning cLogP. The value of the rule is indicated by the grey vertical line, and the shaded area of the chart represents compounds that would fail this rule

**Figure 4.**
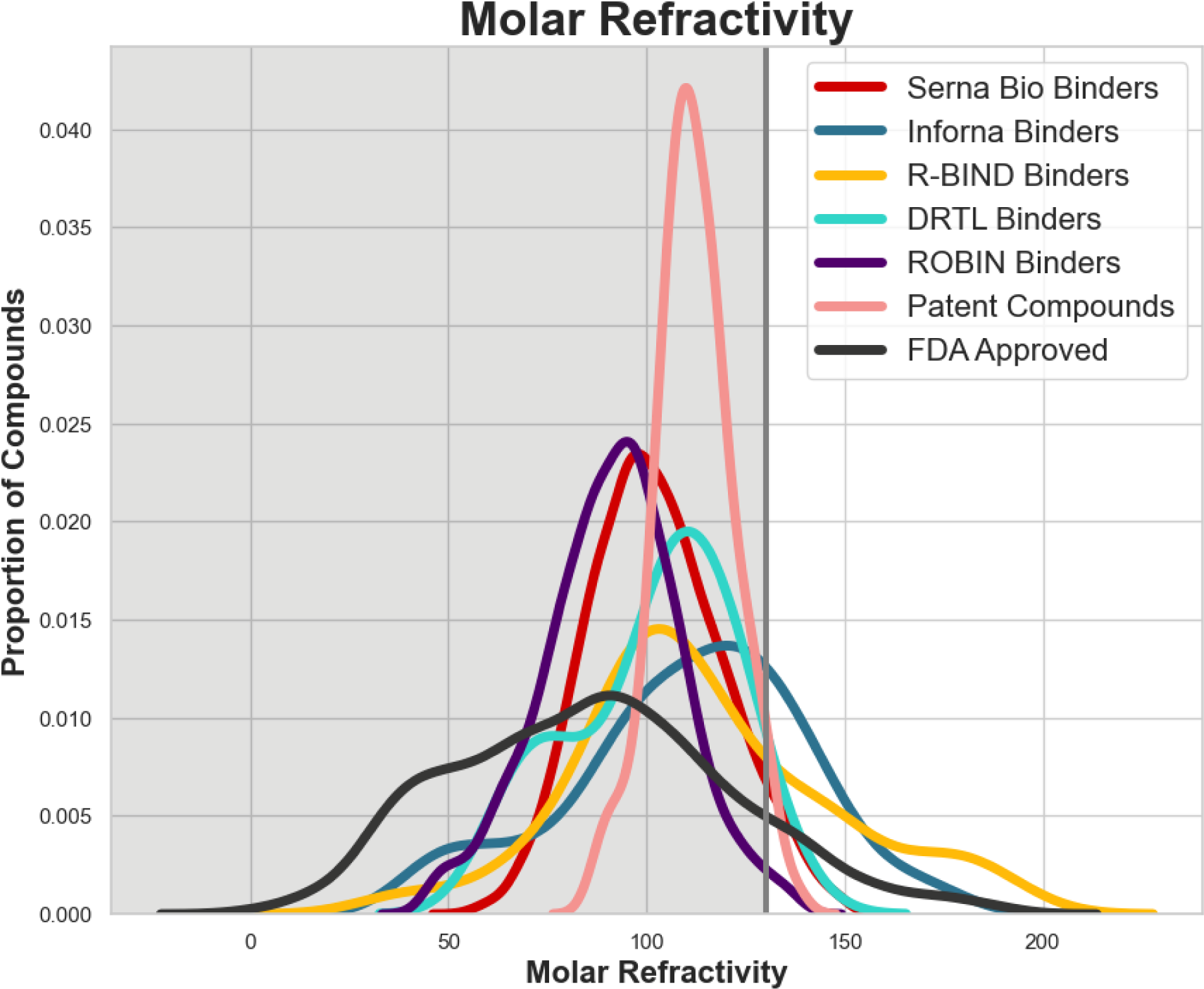
A kernel density estimate distribution plot showing the distribution of Molar Refractivity values for Serna Bio binders (red), ROBIN binders (purple), R-BIND binders (yellow), DRTL binders (turquoise), Patent Compounds (pink), Inforna Binders (blue) and FDA Approved Drugs (grey) with a comparison to the rule we have outlined concerning Molar Refractivity. The value of the rule is indicated by the grey vertical line, and the shaded area of the chart represents compounds that would fail this rule

**Figure 5.**
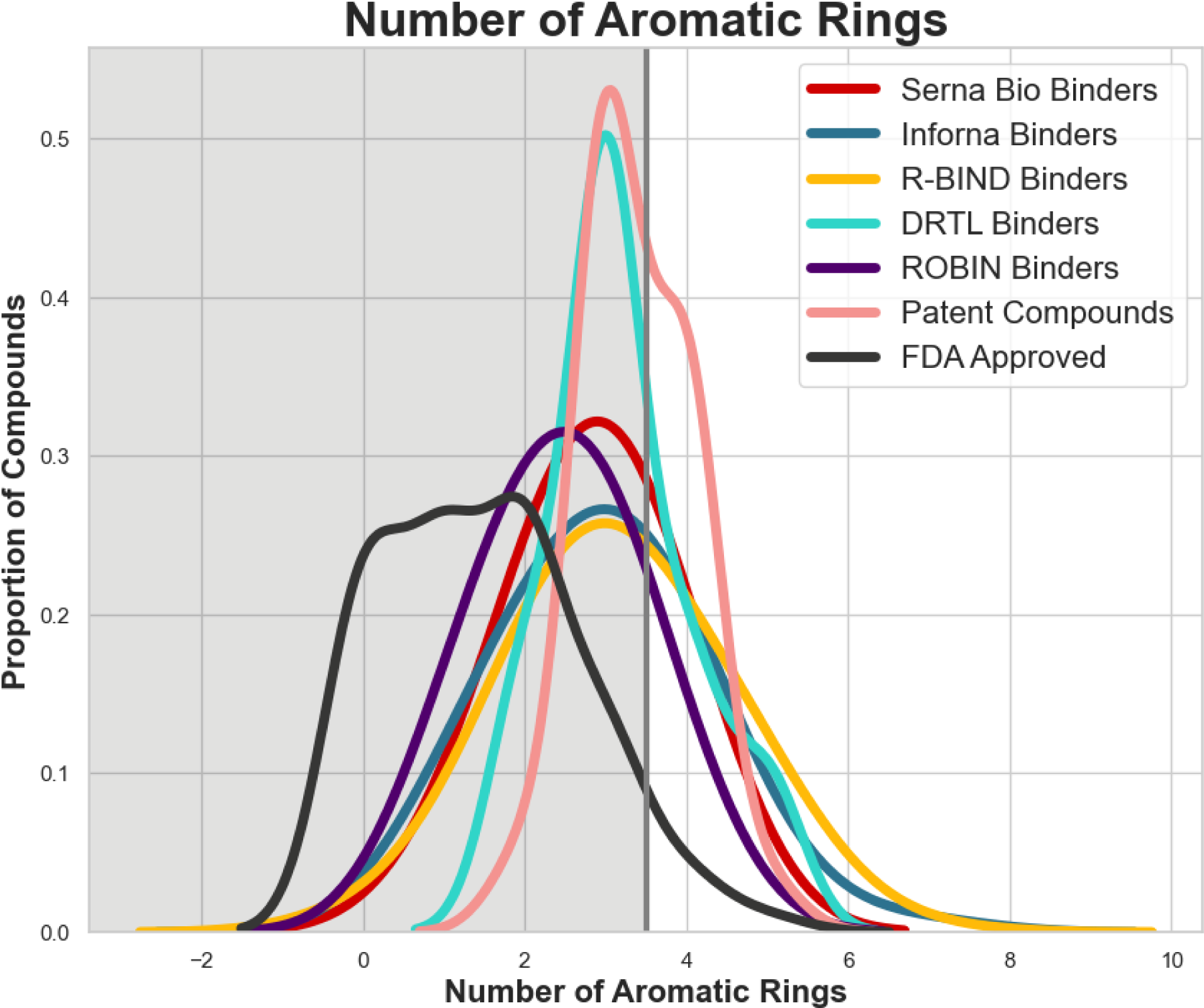
A kernel density estimate distribution plot showing the distribution of the Number of Aromatic Rings in Serna Bio binders (red), ROBIN binders (purple), R-BIND binders (yellow), DRTL binders (turquoise), Patent Compounds (pink), Inforna Binders (blue) and FDA Approved Drugs (grey) with a comparison to the rule we have outlined concerning the Number of Aromatic Rings. The value of the rule is indicated by the grey vertical line, and the shaded area of the chart represents compounds that would fail this rule

**Figure 6.**
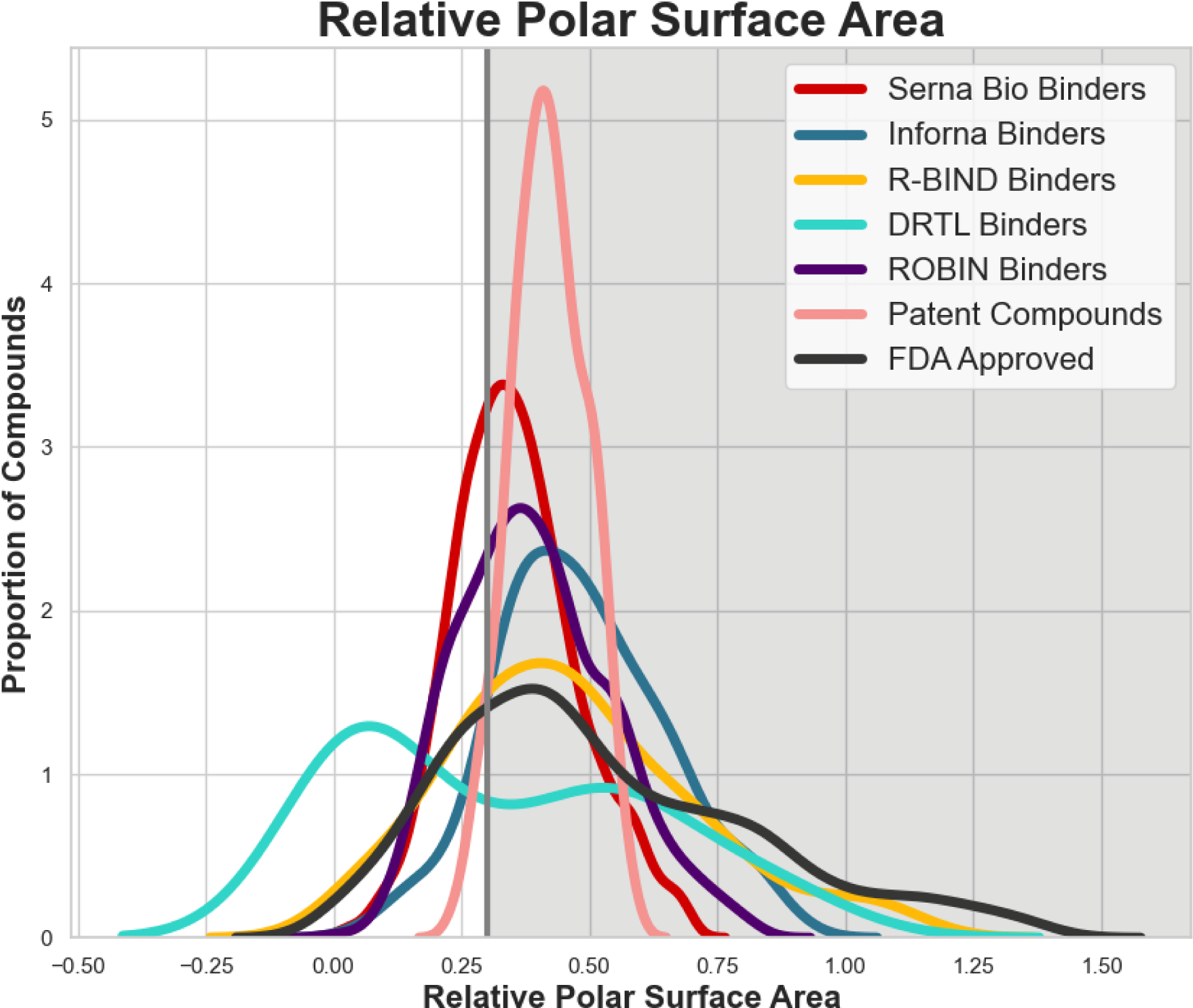
A kernel density estimate distribution plot showing the distribution of Relative Polar Surface Area values for Serna Bio binders (red), ROBIN binders (purple), R-BIND binders (yellow), DRTL binders (turquoise), Patent Compounds (pink), Inforna Binders (blue) and FDA Approved Drugs (grey) with a comparison to the rule we have outlined concerning Relative Polar Surface Area. The value of the rule is indicated by the grey vertical line, and the shaded area of the chart represents compounds that would fail this rule

We emphasize that, like Lipinski’s Rule of Five for druglike compounds,^28^ our STaR rules-of-thumb describe physicochemical tendencies of RNA binders rather than strict, dichotomous cutoffs for these properties. Proven RNA binders need not pass any or all of the STaR rules, though compounds which do are more likely to bind RNA. That said, the idea of RNA binders being more lipophilic, more polarizable, having more aromatic rings and having higher relative polar surface area does align with the following published research: R-BIND noted G-quadruplex binding ligands have a high number of aromatic rings.^16^ The RNA binders found as part of the DRTL study tended to have relatively high LogP.^18^ Analysis conducted as part of the ROBIN study suggested that RNA binders are found to have higher values of chemical descriptors describing aromatic ring systems.^17^ Work based on the Inforna data set identified important chemical scaffolds, several of which contain aromatic systems.^14^ Regarding the patent data, we assume that these molecules have been protected due to evidence of their ability to bind and/or have a functional effect biologically. However, as patent protection can cover a range of potential molecules it is not clear that all have been confirmed as binders. It is encouraging to see that a large percentage (41%) of the patent data set molecules meet the STaR rules.

### FDA-Approved Drugs and the STaR Rules

Our STaR rules for RNA binders can be applied to FDA-approved drugs to illustrate the overlap in physicochemical space between FDA-approved compounds and RNA binders. This was performed using the SelleckChem FDA-approved screening library.^31^ In total 842 of the 2481 compounds in our FDA-approved drug set (∼34%) pass the STaR rules. This aligns well with the total number of Serna Bio and ROBIN compounds that pass. The interaction between the STaR rules and FDA-approved compounds can be further explored by examining Figures 3-6 and the distribution plotted in grey.

To further explore the specific relationship between the STaR rules and RNA binding approved drugs, we calculated the physicochemical properties of Risdiplam^5^ and the RNA binding approved drugs identified by Fang *et al*.^19^ Risdiplam was found to pass the STaR rules, as were 5 of 9 (56%) of the RNA binding approved drugs identified by Fang *et al.* These results further support our conclusions that the STaR rules are generalizable and can be used to narrow the chemical search space for RNA binding approved drug candidates.

The interaction between the FDA-approved drugs and STaR rules confirms that the physicochemical space we have identified is not outside druglike chemical space. In fact, it could be considered that some of the FDA-approved drugs that pass the STaR rules are viable candidates for exploration in the search for RNA binders. This presents potentially interesting questions regarding the consequences that might arise from FDA-approved compounds interacting with RNAs, and how this might be impacting the effect they are having *in vitro* and *in vivo*.^19^

Furthermore, if we consider the drug-likeness of the RNA binders we have identified in the Serna Bio dataset, more than 97% of them pass Lipinski’s rules^28^ - providing robust evidence that these compounds can both bind to RNA and be good starting points for drug discovery efforts. When considering Lipinski’s rules and drug-likeness, one rule that particularly stands out is the rule concerning cLogP, as Lipinski’s rules indicate that compounds having cLogP > 5.0 are less likely to be druglike. As our rule sets as cLogP ≥ 1.5, this leaves a considerable amount of space for compounds to pass both the STaR rules and Lipinski’s rule regarding cLogP, although it should be noted that passing the STaR rules does not imply that a compound is orally bioavailable as Lipinski’s rule does.

## Conclusions

In this work, we have used a statistical analysis of public and proprietary RNA binding data to devise the STaR rules - physicochemical rules of thumb that can be used to define areas of chemical space enriched for RNA binders inspired by the idea of the Rule of Five. The STaR rules can be summarized as follows:

- As a rule of thumb, RNA binders have at least two of the following four properties:

○ cLogP ≥ 1.5
○ Molar Refractivity ≥ 130
○ Number of Aromatic Rings ≥ 4
○ Relative Polar Surface Area ≤ 0.30

The STaR rules were defined using Serna Bio RNA binding data consisting of more than 5,000 RNA binders in a dataset of more than 181,000 small molecules and were found to also be applicable to the R-BIND, DRTL and ROBIN publicly available datasets. The STaR rules for RNA binders use some different chemical properties than those previously developed for drug-likeness, but can provide guidance to chemists working on small molecule RNA drug discovery. The STaR rules are easily and rapidly calculable, from 2D chemical structures using open-source computational packages such as RDKit, Open Babel and Mordred,^23,32,33^ and can be gleaned from chemical structures directly (in the case of the number of aromatic rings) or potentially intuited (in the case of cLogP, molar refractivity or relative polar surface area). Furthermore, the STaR rules provide the possibility for physicochemistry-derived RNA targeting chemical libraries to enable the discovery of new RNA binders at an increased hit rate. In addition, analysis of FDA-approved drugs revealed our STaR ruleset to be compatible with druglike chemical space, and the Serna Bio RNA binders were found to be druglike according to the Rule of Five at a rate of more than 97%.

The thorough physicochemical analysis introduced in this work enables a greater understanding of the chemistry of RNA binders and provides clear evidence that the kinds of chemical characteristics that drive RNA binding are not exclusionary to approved drug molecules. The findings are currently limited to RNA binders due to a lack of availability of functional RNA cellular response data, but the techniques developed could be applied to this kind of data as well, allowing for the derivation of rules of thumb for functional RNA binding in the near future. Knowledge of the chemical drivers of interactions between small molecules and RNA, and functional responses to these interactions help lay the foundation for more drug discovery campaigns where an RNA molecule is the primary target.

## Materials and Methods

### Datasets

Publicly available RNA Binding data was obtained from R-BIND (https://rbind.chem.duke.edu/),^16^ the Supplementary Material for DRTL (https://www.biorxiv.org/content/10.1101/2023.07.31.551350v1)^18^ and the GitHub repo accompanying ROBIN (https://github.com/ky66/ROBIN/tree/main).^17^ These datasets all contain SMILES strings, or compound images that were converted to SMILES strings manually. These SMILES strings were desalted and canonicalized using RDKit^32^ and duplicate canonicalized SMILES strings were removed.

The Inforna data set was kindly provided by the original authors.^13,14^ The data set contained the SMILES strings of each molecule. The SMILES strings were desalted and canonicalized using RDKit^32^ and any duplicate canonicalized SMILES strings were removed.

The patent data set was kindly provided by colleagues at AbbVie. The data set was provided with SMILES strings and underwent the same processing of desalted and canonicalized using RDKit^32^ and any duplicate SMILES strings were removed

Serna Bio data was collected through the screening of commercially available small molecule compound collections totalling more than 181,000 unique small molecules against RNA targets synthesised by Dharmacon, IDT or GenScript. Binding data was obtained using the automated ligand identification system (ALIS)^24^ at PureHoney Technologies/Momentum Biotechnologies as well as using the self-assembled monolayer desorption ionization-affinity selection-mass spectrometry (SAMDI-ASMS)^25^ system at SAMDI Tech. Both assays are examples of ASMS technology.

In the case of ALIS, an RNA target (5μM of each) was incubated for 30 min at room temperature with compounds in pools of 320, 300 or 5 compounds of known mass (1μM each, final DMSO concentration of 3%). Resulting RNA-small molecule complexes were stabilized at 4 °C and separated from unbound compounds using size exclusion chromatography (column: 2.1mm I.D. × 50mm, PolyHydroxyethyl A, 60 Å, (PolyLC, Columbia MD); solvent: 700mM ammonium acetate, pH 8; flow rate: 0.3mL/min at 4 °C). Isolated complexes were then delivered to a reverse phase chromatography column (Targa C18, 0.5mm I.D. × 50mm, 5 μm packing material (Higgins Analytical, Mountain View, CA) running at a linear gradient of 0-90% acetonitrile/0.2% formic acid in 2.5 min (total run time of 4 min including column equilibration) at 20μL/min and 60 °C) to dissociate, desalt and separate small molecules for subsequent detection using electrospray ionization-based mass spectrometry (Thermo Scientific™Exactive™ Orbitrap high-resolution, accurate-mass mass spectrometer detector operating at 100,000 resolution with a mass accuracy of <5 ppm and a scan rate of 1 Hz). Binders were selected using the following criteria: uniqueness (absent from the blank control), ASMS score > 75 (depends on mass accuracy and isotopic distribution), inspection of peak area and S/N ratio.

In the case of SAMDI-ASMS, a biotinylated RNA target (0.4μM of each) was incubated for 60 min at room temperature with pools of 8 or individual compounds of known mass (15μM each, final DMSO concentration of 1.2%). Subsequently, the mixtures were transferred to SAMDI biochip arrays functionalized with neutravidin and incubated for 60 min in a humidified chamber to allow for immobilization of RNA-small molecule complexes. The arrays were rapidly washed with ultra-pure water and dried with compressed air prior to matrix (alpha-cyano cinnamic acid) addition and analysis by matrix assisted laser desorption ionization (MALDI) mass spectrometry (AB Sciex TOF-TOF 5800 System (AB Sciex, Framingham, MA), reflector positive mode, 400 shots/spot, 400 Hz laser frequency, bin size of 1 ns, and detector voltage multiplier of 0.48. A mass window of m/z 230 to m/z 900 was used and a mass threshold of m/z 0.5 applied for peak identification). In order to minimize matrix-related MALDI signal variability, the area under the curve (AUC) of each peak that corresponds to the compound’s mass ID was normalized to the sum of the compound’s AUC and the AUC of the the internal comparator molecule (m/z 335.2), generating a relative signal value (RSV). In addition, to account for compound selectivity, a signal to background (S/B) value was calculated for each compound by dividing its RSV generated in the presence of the RNA by the RSV generated in the absence of the target. Binders were then selected by setting thresholds on RSV (>1 standard deviation above the average RSV for all compounds for a given target) and S/B ratio (>2 times the average S/B for all compounds for a given target). Compounds that passed the set thresholds underwent additional QC steps where their MS spectra were evaluated with respect to uniqueness, signal intensity and isotopic distribution pattern.

As a comparison dataset, an example of FDA-approved compounds is represented by the SelleckChem FDA-approved screening library (https://www.selleckchem.com/screening/fda-approved-drug-library.html)^31^ which contains 2498 unique, desalted, canonicalized SMILES strings.

In the Serna Bio and ROBIN datasets, compounds were classified as RNA binders if they showed Binding activity against any of the tested RNA targets in that dataset. Compounds were classified as non-binders if they were non-binders against every tested RNA target. For statistical comparison between RNA binders and non-binders for R-BIND and DRTL, where the datasets contain no compounds labelled as non-binders, the non-binders from the Serna Bio and ROBIN datasets were used to fill this role.

### Analysis of FDA-approved compounds as RNA binders

Statistical significance p-value calculations to compare the hit rates of different classes of FDA-approved drugs to a diverse chemical library were performed using Fischer’s exact test^27^ implemented using SciPy version 1.11.3.^34^

### Physicochemical Calculations and Statistical Analysis

Physicochemical properties were calculated using RDKit version 2022.09.5^32^ via Python version 3.10.4.^35^ Molecules were presented as canonicalized SMILES strings to RDKit for property calculations. Physicochemical profiles of RNA binders and non-binders were then made comparatively using box and whisker plots and p-value calculations using Mood’s median test^29^ with Benjamini-Hochberg correction^30^ implemented using SciPy version 1.11.3.^34^ In the case of the comparisons made, a physicochemical property change was considered to be statistically significant where both i) the Median value changes between the distributions of RNA binders and non-binders, and ii) the p-value was < 0.01. These conditions were used to be more stringent in identifying physicochemical properties that are changing between RNA binders and non-binders in comparison to using a significance level of 0.05, as the large number of data points involved can lead to distributions that look very similar being flagged as statistically significantly different.

### Identification of the STaR Rules

Following the identification of important physicochemical properties using the calculated p-values, a statistical grid search was implemented to identify where physicochemical rule thresholds could be drawn. For each physicochemical property being considered in the STaR rules, testing thresholds were established by breaking the Serna Bio dataset into bins of equal size and taking the values separating those bins. Five values were identified for each property and then all possible combinations of those properties were tested in turn in a complete grid search (see the Supplementary Material for tested thresholds, Table S1). This search included a provision for compounds failing different numbers of components of the rules while still being able to pass overall (for example, a case where a compound can meet two of the four criteria and still be considered a pass - as in the final identified STaR rules). This idea is inspired by Lipinski’s Rule of Five in which compounds must pass three of the four physicochemical bounds to be considered druglike, but do not have to pass all four.^28^ Where RNA binders were found to have a higher median physicochemical value compared to non-binders, a property was required to be greater than or equal to its threshold, and vice-versa for lower median physicochemical values. In the case of each proposed ruleset, the number of compounds, and the number of RNA binders in the Serna Bio dataset passing the ruleset were identified, as well as the Compound Hit Rate defined as

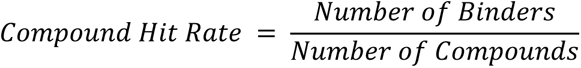

Compound Hit Rate acts as a measure of the proportion of passing compounds that are RNA binders.

To establish a ruleset that is generalizable to a notable number of RNA binders, any ruleset that did not pass 50% of the RNA binders in the Serna Bio dataset was excluded. The highest Compound Hit Rate of the remaining rulesets was identified as the best starting point for the rules of thumb we wished to obtain. To simplify the values to appropriate rules of thumb along the line of the Rule of Five, the physicochemical thresholds in the best ruleset were rounded and retested on the Serna Bio dataset to ensure they were still enriched for RNA binders. The physicochemical thresholds in the STaR rules were rounded as follows:

- The cLogP threshold was rounded from 1.81 to 1.5, as rounding to 2 would have resulted in RNA binders with cLogP between 1.81 and 2 falling outside the threshold and being rejected by the STaR rules
- The molar refractivity threshold was rounded from 136.70 to 130, as rounding to 140 would have resulted in RNA binders with molar refractivity between 136.70 and 140 falling outside the threshold and being rejected by the STaR rules
- The aromatic rings threshold was rounded from 3.43 to 4, as this descriptor is always an integer and the same molecules will be selected
- The relative polar surface area threshold was rounded from 0.3021 to 0.30 as rounding to the nearest two significant figures. Rounding to 0.40, 0.50 or 0.25 all resulted in a notable decrease in Compound Hit Rate for Serna Bio compounds passing the rule

These STaR rules were then applied to R-BIND, DRTL, ROBIN, Inforna and Patent compounds for external validation and FDA-approved compounds to examine the drug-likeness of compounds passing them.

## Supporting information

Physicochemical Principles Driving Small Molecule Binding to RNA - Supplementary Material

Physicochemical Principles Driving Small Molecule Binding to RNA - Supplementary Material Sheets

## Supplementary Material

The SMILES strings of compounds in the patent dataset, the full list of calculated rulesets from which the STaR rules are defined, a further comparison of the STaR physicochemical properties for Serna Bio and ROBIN binders and non-binders and the STaR rule grid search property thresholds are all provided in the Supplementary Material.

## Abbreviations

ASA: Accessible surface area
ASMS: Affinity selection-mass spectrometry
CC-DDR: cell cycle and DNA damage repair
cLogP: calculated LogP
DRTL: Duke RNA targeted library
PBF over MW: Plane of best fit over molecular weight
R-BIND: RNA-targeted bioactive ligand database
Relative PSA: Relative polar surface area
ROBIN: Repository of binders to nucleic acids
STaR: Small molecule targeting RNA
TPSA: Topological polar surface area

