## Supplementary material for "Physicochemical Principles Driving Small Molecule Binding to RNA": Physicochemical Principles Driving Small Molecule Binding to RNA - Supplementary Material

*[1]Serna Bio*
*[2]Department of Molecular Genetics, Groningen Biomolecular Sciences and Biotechnology Institute (GBB), University of Groningen, Groningen, the Netherlands*
*[3]AbbVie, 1 North Waukegan Road, North Chicago, IL 60064, United States of America*

*

**Contents:**

Patent Compounds Dataset S2
Full Set of Calculated Results S3
Physicochemical KDE Distribution Plots of Binders and Non-Binders S4-S7
Table of Tested Physicochemical Thresholds S8

### Patent Compounds Dataset

**Tab 1.0 - Patent Compounds Dataset** in Physicochemical Principles Driving Small Molecule Binding to RNA - Supplementary Material Sheets.xlsx

### Full Set of Calculated Rulesets

**Tab 2.0 - Full Set of Calculated Rulesets** in Physicochemical Principles Driving Small Molecule Binding to RNA - Supplementary Material Sheets.xlsx

### Physicochemical Kernel Density Estimate Distribution Plots of Binders and Non-Binders

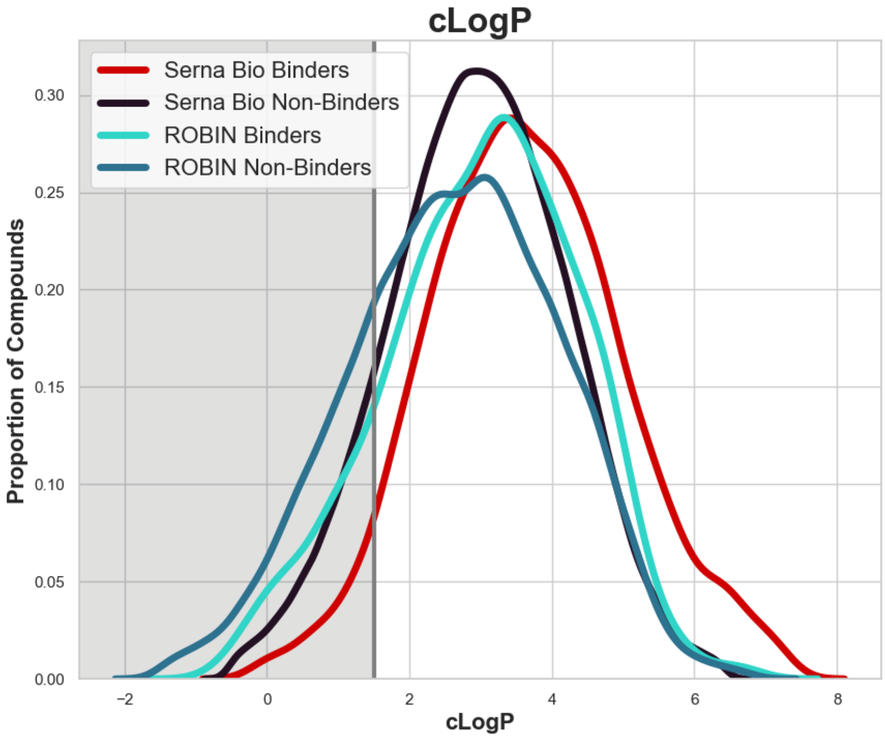


*Figure S1 - A kernel density estimate distribution plot showing the distribution of cLogP values for Serna Bio binders (red), Serna Bio non-binders (purple), ROBIN binders (turquoise), and ROBIN non-binders (blue) with a comparison to the rule we have outlined concerning cLogP. The value of the rule is indicated by the grey vertical line, and the shaded area of the chart represents compounds that would fail this rule*

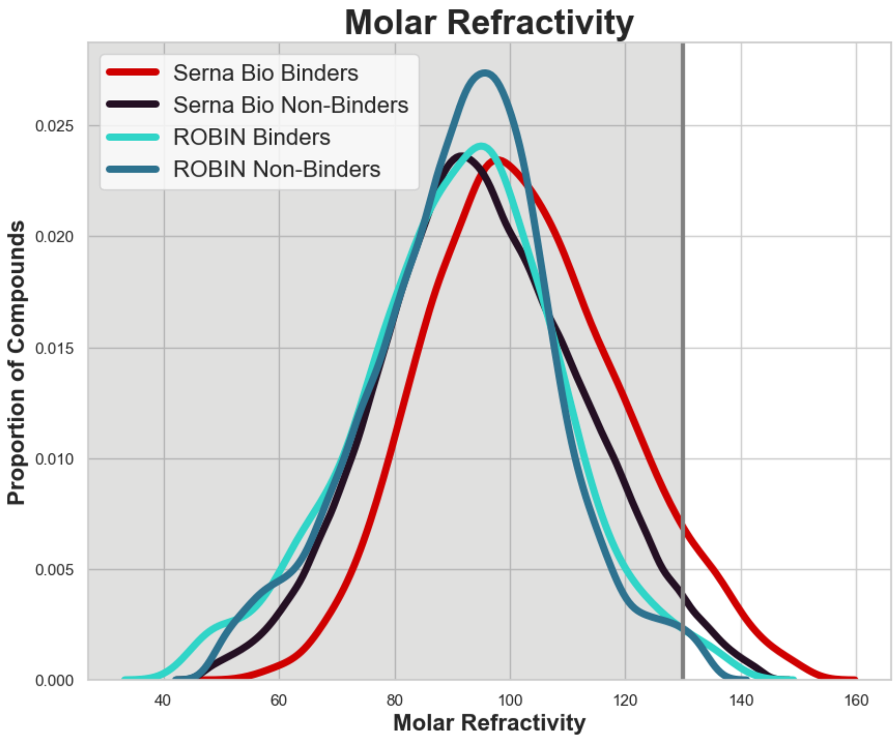


*Figure S2 - A kernel density estimate distribution plot showing the distribution of Molar Refractivity values for Serna Bio binders (red), Serna Bio non-binders (purple), ROBIN binders (turquoise), and ROBIN non-binders (blue) with a comparison to the rule we have outlined concerning Molar Refractivity. The value of the rule is indicated by the grey vertical line, and the shaded area of the chart represents compounds that would fail this rule*

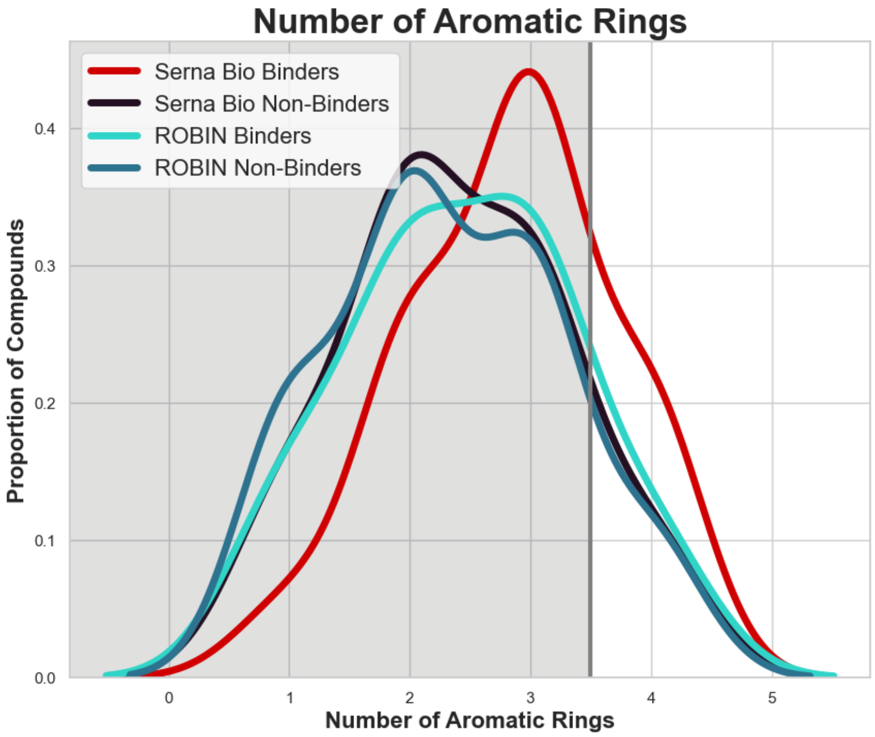

*Figure S3 - A kernel density estimate distribution plot showing the distribution of the Number of Aromatic Rings in Serna Bio binders (red), Serna Bio non-binders (purple), ROBIN binders (turquoise), and ROBIN non-binders (blue) with a comparison to the rule we have outlined concerning the Number of Aromatic Rings. The value of the rule is indicated by the grey vertical line, and the shaded area of the chart represents compounds that would fail this rule*

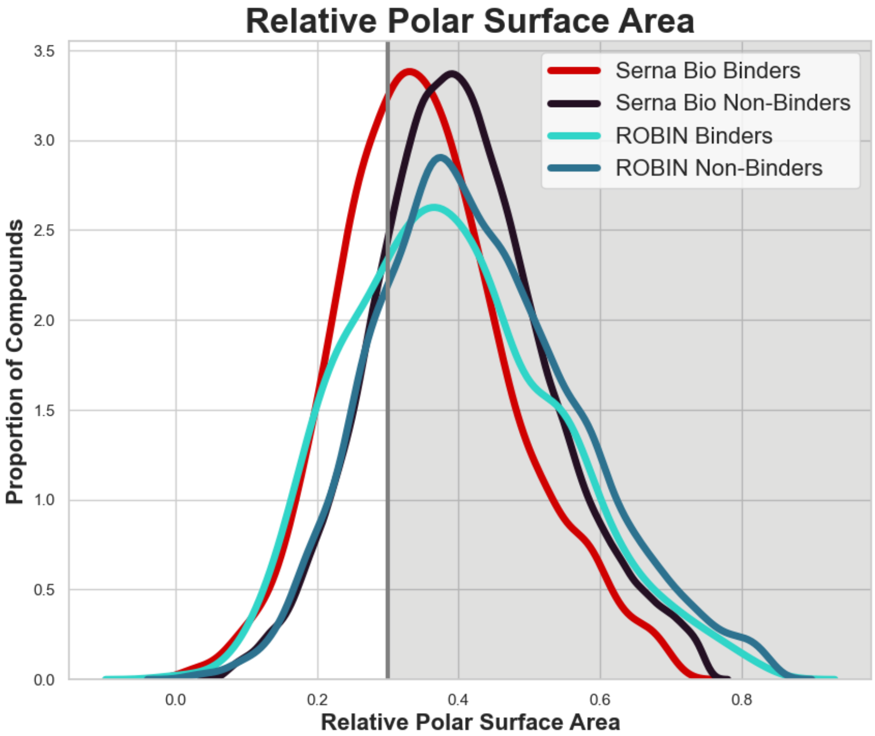


*Figure S4 - A kernel density estimate distribution plot showing the distribution of Relative Polar Surface Area values for Serna Bio binders (red), Serna Bio non-binders (purple), ROBIN binders (turquoise), and ROBIN non-binders (blue) with a comparison to the rule we have outlined concerning Relative Polar Surface Area. The value of the rule is indicated by the grey vertical line, and the shaded area of the chart represents compounds that would fail this rule*

### Table of Tested Physicochemical Thresholds

| Threshold Number | cLogP | Molar Refractivity | Number of Aromatic Rings | Relative Polar Surface Area |
|---|---|---|---|---|
| Threshold 1 | -14.23 | 79.41 | 1.14 | 0.3021 |
| Threshold 2 | -8.89 | 136.70 | 2.29 | 0.6041 |
| Threshold 3 | -3.54 | 193.99 | 3.43 | 0.9062 |
| Threshold 4 | 1.81 | 251.28 | 4.57 | 1.2083 |
| Threshold 5 | 7.16 | 308.58 | 5.71 | 1.5103 |

*Table S1 - The thresholds for each physicochemical property tested in the grid search to define the STaR rules*
